# Humans share acoustic preferences with other animals

**DOI:** 10.1101/2025.06.26.661759

**Authors:** Logan S. James, Sarah C. Woolley, Jon T. Sakata, Courtney B. Hilton, Michael J. Ryan, Samuel A. Mehr

## Abstract

Many animals produce sounds during courtship and receivers prefer some sounds over others. Shared ancestry and convergent evolution may generate similarities in preference across species and could underlie Darwin’s conjecture that some animals “have nearly the same taste for the beautiful as we have”. Here, we show that humans share acoustic preferences with a wide range of animals, that the strength of human preferences correlates with that in other animals, and that human responses are quicker when in agreement with animals. Furthermore, we found greatest agreement in preference for adorned, evolutionarily ancestral, and lower frequency sounds. Humans’ music listening experience was associated with preferences. These results are consistent with theories arguing that biases in sensory and cognitive processing sculpt acoustic preferences and confirm Darwin’s century-old hunch about the conservation of aesthetics in nature.

## Main Text

Many animals produce signals to attract mates. These signals span sensory modalities from visual color patterns and movements to acoustic songs and calls to olfactory plumes, and are produced throughout the animal kingdom, including arthropods, molluscs, and all vertebrate classes (*1*).

The receivers of mating signals generally exhibit preferences for the signals of conspecifics relative to those of other species (*1, 2*), and mating with conspecifics usually results in more viable offspring (*3, 4*). Moreover, mating signals typically vary within a species, and receivers often exhibit preferences for some signal variants over others, which may result from sensory biases (*5–15*), adaptive pressures (*16–20*), or both.

Preferences emerge from an interaction between the signal’s attributes and the receiver’s sensory system. For example, stimuli that evoke greater stimulation of the sensory system are often preferred over stimuli that are less stimulating (*14, 21*). There is substantial conservation in the organization of sensory systems across species (e.g., *22*), which may explain why the dazzling colors of butterflies, aromas of flowers, and songs of birds are attractive not just to their intended receivers, but to humans as well.

Darwin suggested that some animals “have nearly the same taste for the beautiful as we have” (*23, 24*). Here, we report a citizen-science experiment testing this hypothesis.

## Results

We gathered 110 pairs of sounds produced by 16 non-human animal species (hereafter, animals), recorded in prior research (*25–50*), and played them to humans recruited globally online (*n* = 4,196 participants, geographic distribution in Fig. S1) in a gamified experiment (*51*). Participants rated which of the two sounds in a pair they “liked more” (see Methods and Fig. S2; *n* = 48,567 responses).

For each stimulus pair, the animals from which the sounds originated are known to display a preference for one of the two sounds (hereafter, the more-attractive sound). For example, male túngara frogs can produce a simple call or a complex one that includes an acoustic adornment; female frogs choose a complex call over a simple call approximately 84% of the time, a 5:1 preference (*41*). The strength of animals’ preferences varied (range: 55%-93%; see Methods and Table S1), enabling tests of the degree to which human preferences were similar to those of other species.

There were three principal findings. First, human preferences for animal sounds correlated with the preferences of the animals themselves: the percent of humans that selected the more-attractive sound correlated positively with the animal’s strength of preference (LMM: F_1,34.9_ = 6.2, *p* = 0.02; Fig. 1A).

**Fig. 1.**
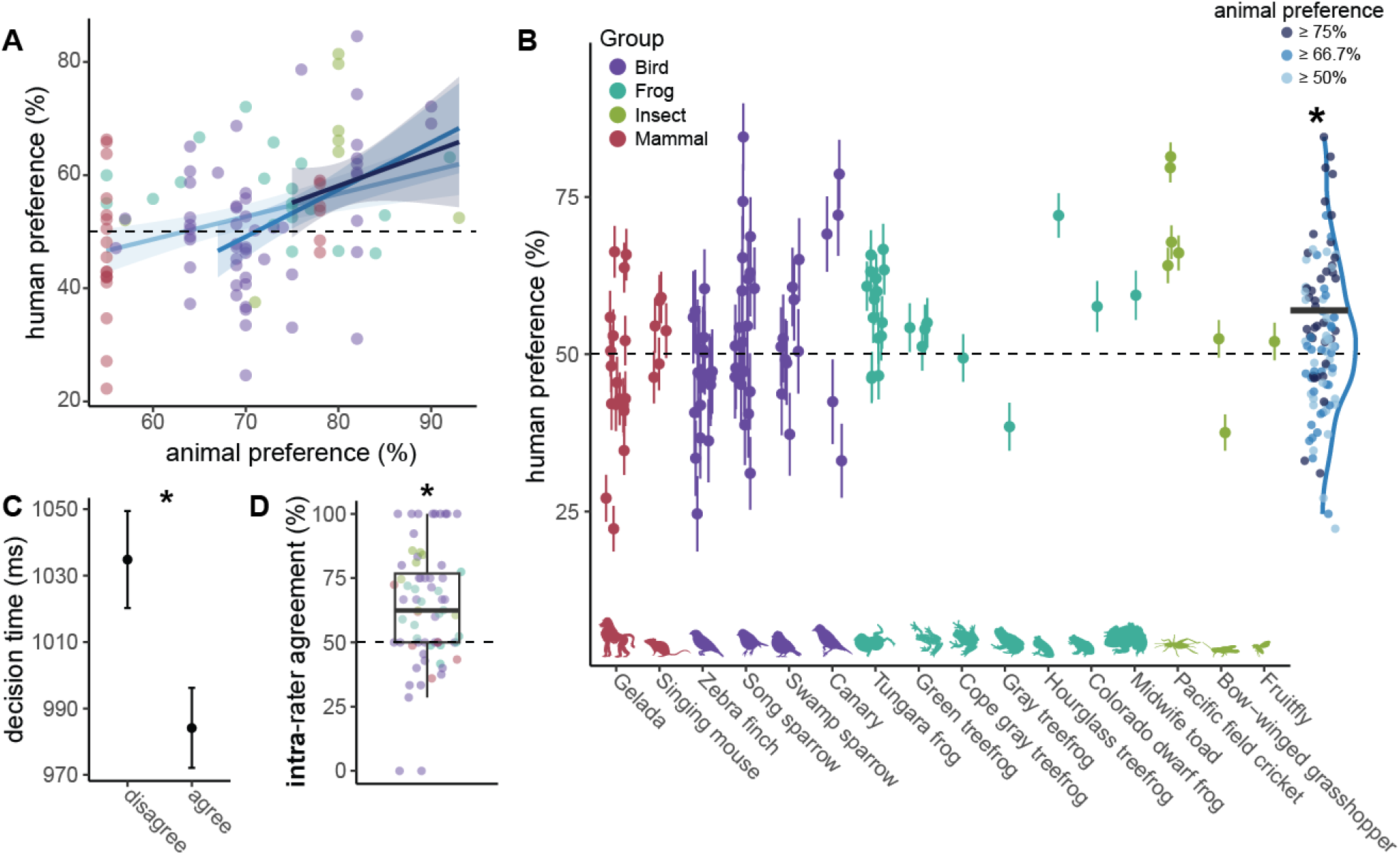
Humans share acoustic preferences with other animals. (**A**) The scatterplot shows the correlation between the strength of the animal’s preference for the more-attractive sound in a stimulus pair (*x*-axis) and the percent of humans that preferred the more-attractive sound (*y*-axis). Each dot represents a stimulus pair. Trendlines show simple linear correlations (shading for standard error) when analyzing all stimuli (light blue); stimuli with at least a 2:1 animal preference (medium blue); or stimuli with at least a 3:1 animal preference (dark blue). (**B**) Humans chose the more-attractive sound above chance. In the main plot, the dots each depict the percent of humans that preferred the more-attractive sound from a given species; the vertical lines represent 95% confidence intervals; and the horizontal dotted line represents chance (50%). To the right, each dot represents the mean level of human preferences for a stimulus pair, with color indicating the strength of preference in animals. The solid blue line represents a kernel density estimation and horizontal bar denotes the mean across stimulus pairs with at least a 2:1 preference in animals. (**C**) Decision-making time was significantly shorter for trials when the participant selected the more-attractive stimulus; the dots depict the mean and the error bars depict 95% confidence intervals, collapsing across all trials (back-transformed from z-scores for visualization). (**D**) When participants heard the same stimulus pair more than once, their preferences were maintained, on average. Each dot depicts the mean intra-rater agreement for a stimulus pair, and the boxplot depicts the median, interquartile range, and 1.5xIQR. Across all panels, the colors correspond to large phylogenetic group. ^∗^*p* < 0.05

Second, agreement between human and animal preferences was higher when the animals’ preferences were more robust. When animals exhibited moderately strong preferences (at least 2:1 odds, or 67% preference; see Methods), humans were significantly more likely than chance to choose the more-attractive sound (GLMM: mean 56.4% agreement, 95% CI: 55.8%-57.0%; *z* = 2.3, *p* = 0.02; Fig. 1B). This effect was slightly weaker when including all stimuli (i.e., including those with subtle animal preferences; 54.0% agreement: *z* = 1.9, *p* = 0.05), and stronger with a more stringent cutoff (at least 3:1 odds: 59.5% agreement; *z* = 3.4, *p* < 0.01). All subsequent analyses use the 2:1 animal preference dataset by default. Moreover, we found no significant differences in agreement across the large clades of animals present in the study (birds, mammals, frogs, or insects; 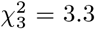, *p* = 0.35).

Third, shared preferences across humans and animals were supported by two additional measures. Humans answered 51 ms faster, on average, when choosing the more-attractive sound than the less-attractive one (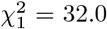, *p* < 0.01; Fig. 1C). Their responses were also internally reliable: in a subset of trials, we played participants the same stimulus pair twice and found their preference was maintained across both presentations at a rate higher than chance (63% were the same choice, on average; *t*_69_ = 5.0, *p* < 0.01; Fig. 1D).

We next tested the degree to which specific properties of animal sounds are predictive of human preferences. The stimuli studied here derived from three classes of experiments, each originally designed to test whether a specific characteristic was predictive of preferences in animals. In planned analyses, we asked whether humans similarly expressed such preferences.

In the first class of experiments, researchers experimentally manipulated the sounds of animals (Table S1). For example, in three pairs of frog calls (*38, 44*) researchers measured frogs’ preferences for frequency-manipulated calls. Our human participants agreed with the frogs (preferring lower frequency calls; GLMM: *z* = 3.9, *p* < 0.01). Similarly, both humans and animals preferred sounds with acoustic adornments, such as “trills”, “clicks”, and “chucks” (*z* = 4.0, *p* < 0.01). There were no significant effects of agreement for stimuli distinguished primarily by rate or amplitude modulation (Fig. 2A).

**Fig. 2.**
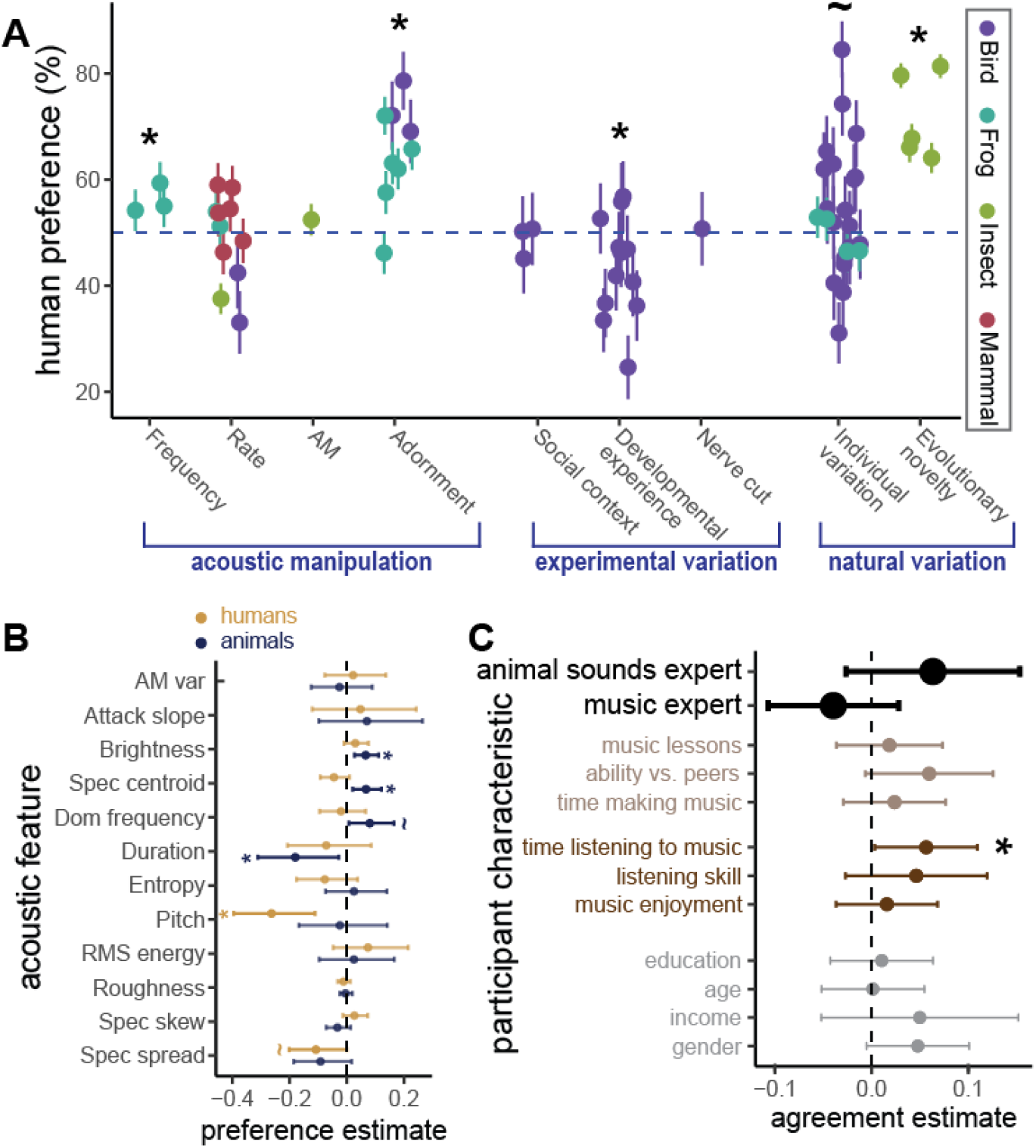
Influence of acoustic traits and participant characteristics on human-animal agreement. (**A**) In some cases, variation within stimulus pairs predicted human preferences. There were three categories of variation: acoustic manipulations; experimental variation, where the animals producing the sounds differed on the basis of a manipulation; or natural variation, where the sounds varied without experimental input. Dots denote the mean human preference for the more-attractive sound in each stimulus pair and the vertical lines denote 95% confidence intervals. (**B**) Dots denote model estimates (*z*-scores) and horizontal lines denote 95% confidence intervals for each acoustic feature on human (orange) or animal (blue) preferences. (**C**) Dots denote estimates from models asking whether a given participant characteristic predicted human agreement with other animals, and the horizontal lines denote 95% confidence intervals (values binarized for visualization). The two large black dots denote planned analyses and smaller dots indicate exploratory analyses relating to music production (light brown), perception (dark brown), or demographics (grey). ^∗^*p* < 0.05, ~*p* < 0.10

In the second class of experiments, researchers studied animals across experimental conditions (e.g., varied social contexts) and tested preferences for the sounds they produced. Surprisingly, humans disagreed with animals for stimuli that varied in the developmental experience of the signaler (*z* = −2.5, *p* = 0.01). Specifically, humans prefer zebra finch songs from males reared without a tutor (“isolate songs”) over songs from birds that learned their song. We found no effect of agreement for songs produced in different social contexts or songs produced with or without a nerve cut (Fig. 2A).

In the third class of experiments, researchers recorded the natural variation in sounds across individuals, and tested animals’ preferences for them. Humans shared a preference for sounds that had less evolutionary novelty; both humans and crickets preferred ancestral “chirps” over novel “purrs” (*z* = 7.3, *p* < 0.01; Fig. 2A). We also observed a non-significant trend toward agreement for general individual differences in acoustic structure (which may signal benefits to the receiver; *z* = 1.7, *p* = 0.08; Fig. 2A).

In exploratory analyses, we also asked whether any single acoustic feature could predict which stimulus was more attractive to humans and/or other animals across all stimuli. Consistent with the idea that preferences arise from complex interactions of multiple cues (*52*), we did not find that a single acoustic feature predicted the behavior of both humans and animals. Animals were more likely to prefer the stimulus with greater brightness, higher spectral centroid, or shorter duration (Fig. 2B; Table S3), whereas humans were more likely to prefer the stimulus with lower pitch (Figs. 2B, S3; Table S3).

Last, we tested whether characteristics of the participants were predictive of their agreement with animals. We predicted that prior experience with animal sounds and musical expertise would predict human preferences given that experience can shape auditory perception and preferences (*53–56*).

Neither prediction was supported. Humans with experience identifying animals by their sounds, such as birders (“animal experts”; *n* = 373, ~9% of participants), and expert musicians (*n* = 730, ~19% of participants) exhibited similar degrees of agreement as non-experts (GLMMs: animal experts: 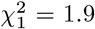, *p* = 0.17; music experts: 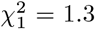, *p* = 0.25; Fig. 2C). The only significant predictor of agreement was time listening to music, with those reporting more time listening to music per day agreeing with animals more (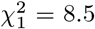, *p* < 0.01; Figs. 2C, S1; Table S3).

In exploratory analyses, we asked whether demographic variables (e.g., age, gender) predict preference; gender is of particular interest given that preferences in male animals can be at least as strong as those in females (*57, 58*). While no demographic variable predicted human preference (Fig. 2C; Table S3), one result was intriguing: analysis of the full dataset (i.e., including the stimuli with weak animal preferences), revealed a small, statistically significant effect of participant gender, where male participants agreed with animals more than female participants. But this result was internally inconsistent (see Methods), and did not hold when limiting the stimulus set to pairs with clear preferences in animals. We therefore suggest caution with interpretation. Finally, human agreement with animals remained significantly above chance (*z* = 2.3, *p* = 0.02), even when adjusting the main analysis for demographic variables (gender, education, age, and income) or time spent listening to music.

## Discussion

We report shared acoustic preferences between humans and other animals, generalizing across insects, frogs, birds, and non-human mammals. The strength of preferences for the more-attractive sound was correlated across humans and animals; humans were more likely than chance to prefer the more-attractive sound; and humans answered more quickly when selecting the more-attractive sound.

These results are consistent with general form-function relationships in acoustic signals across species (*59–65*). For example, sound “roughness” is observed in alarm and distress calls and is perceived as aversive across numerous species (*52, 66–72*). Humans appear to be sensitive to this and other form-function relationships in other species (*73–76*). Moreover, some animals are sensitive to distress calls from other species and many animals respond to heterospecific alarm calls (*77, 78*). While most studies have focused on sounds used in aggressive, aversive, or submissive contexts, our results suggest that cross-species form-function relationships may similarly be used for preference. Such relationships may underlie the perception of attractiveness in our own communicative vocalizations, including speech and music (*52, 79–81*).

There is little evidence about specific features that robustly predict preferred sounds (*52, 66, 80*), consistent with our finding that no single acoustic feature predicted both human and animal preferences across all stimuli. Relatively ambiguous measures of sound (such as “complexity”) are often proposed to explain preferences (*82–86*). For example, flycatchers and starlings prefer individuals with more complex (larger) repertoires (*83, 84*). Humans similarly prefer more complex birdsongs (*87*), but these measures are difficult to generalize across varied species (*88, 89*).

We did find that humans (but not animals) generally preferred the stimulus with lower pitch (*90*), which may relate to human-specific pitch perception (*91*). Overall, there were few similarities across animals and humans in the acoustic characteristics that served as cues to attractiveness — despite their shared overall preferences. Humans’ and animals’ attractiveness inferences likely result from multiple, interacting cues, perhaps reflecting cross-species differences in auditory processing (*22, 91, 92*).

Whereas we did not find an effect of experience with animal sounds [cf. (*55, 93*)] or music production expertise on alignment with animal preferences, we did find that higher levels of music listening experience predicted whether humans agreed with animals. Music training is known to be associated with advantages in auditory processing ability (review: *94*), and at least one study shows an association between auditory discrimination abilities and music listening time (*95*). Perhaps sustained music listening leads to increased attention, or better auditory discrimination ability, which translated to higher concordance with animals [whose vocalizations can share features with music; e.g., (*96, 97*)].

We note that preferences in animals are often subtle, context-dependent, and can vary across individuals and populations (*98*). This may account for the high degree of variation in the strength and direction of human preferences observed across stimuli, and the relatively subtle main effect of agreement between humans and other animals. The overall pattern of convergent evidence from multiple analysis approaches, combined with the increasing robustness when limiting the sample to stimulus pairs for which animals themselves show stronger preferences, suggests a robust main effect.

In sum, these results confirm that a single species can harbor similar preferences to a taxonomically broad range of species. While Darwin’s original idea of shared preferences alluded to the visual coloration of birds, our findings suggest a more expansive shared “taste for the beautiful” (*23, 24*), motivating work across many species and in other modalities (*99*). Finally, these results remind us that much of the beauty we find in nature was intended for receivers other than ourselves.

## Acknowledgments

We thank Elizabeth Adkins-Regan, Jaime Bosch, Katherine Buchanan, Bruce Byers, Morgan Gustison, Albertine Leitão, Stefan Leitner, Stephen Nowicki, Bret Pasch, Susan Peters, Michael Reichert, Michael Ritchie, Ryan Taylor, Robin Tinghitella, Michelle Tomaszycki, Eric-Marie Vallet, and the Macaulay Library for generously providing acoustic stimuli; Jan Simson, Isaac Kinley, and Sophia MacNeil for help with code and data management; and Mila Bertolo, Lidya Yurdum, and the members of the Music Lab for feedback on the design of the experiment and on the manuscript.

## Funding

Smithsonian Institution Postdoctoral Fellowship (LSJ)

Fonds de recherche du Québec – Nature et technologies PR-299652 (SCW, JTS)

US National Institutes of Health DP5OD024566 (SAM)

Royal Society of New Zealand Te Apārangi RDF-UOA2103 (SAM)

## Author contributions

Conceptualization: LSJ, SCW, JTS, MJR, SAM

Methodology: LSJ, JTS, CBH, SAM

Data curation: LSJ, CBH, SAM Investigation: LSJ Visualization: LSJ

Software: LSJ, CBH

Funding acquisition: LSJ, SCW, JTS, SAM

Project administration: SAM

Supervision: SCW, JTS, MJR, SAM

Writing – original draft: LSJ, SCW, JTS, CBH, MJR, SAM

Writing – review & editing: LSJ, SCW, JTS, CBH, MJR, SAM

## Data and materials availability

Data, code, materials, and a reproducible manuscript are available at https://github.com/themusiclab/animal-sounds.

## Supplementary Materials

Materials and Methods

Supplementary Text

Tables S1 to S3

Figures S1 to S3

References 100-105

## Materials and Methods

### Participants

Participants were visitors to the citizen-science website https://themusiclab.org between 2023-09-06 and 2025-09-15 (Fig. S1). Participants were recruited through word of mouth, classroom activities, social media, and targeted newsletters (e.g., Animal Behavior Society). They were not compensated.

We analyzed data from participants who (i) reported age between 18-90; (ii) reported wearing headphones and/or “in a very quiet place”; (iii) reported no hearing impairment; and (iv) reported fluency in English, either as their native or second language. These inclusion criteria were determined before analyzing any data, so could not bias the results in any direction. All participants gave informed consent under ethics protocols approved by the Yale University Human Research Protection Program (protocol 2000033433) and the McGill University Research Ethics Board (initial approval October 24, 2022; protocol 22-09-042).

### Stimuli

We obtained audio files used in a variety of published studies, with the exception of a few stimuli which came from the unpublished collections of colleagues or a public sound library (Macaulay Library; Table S1). In general, audio was requested from scientists by the first author (L.S.J.) based on their studies of conspecific acoustic preferences (i.e., in research where preferences had been measured in receivers of the species that produced the sound). In general, we opted not to use stimuli where the most salient difference between stimuli was in amplitude or duration. All stimuli were produced by male animals, and preference studies were conducted with females as the receivers.

For 14 out of 24 sets of stimuli, stimulus pairs were experimentally manipulated by the researchers. For example, in singing mouse songs, trill rates were experimentally modified digitally. These techniques are common in studies of animal preference, which typically seek to control for precise differences between pairs of sounds. Because we aimed to study the concordance between human preferences and animal preferences, and because animal preferences had been measured using these stimuli in previous work, we treated manipulated stimuli interchangeably with unaltered ones. Table S1 indicates which stimuli were fully naturalistic recordings and which were altered for the preference experiment(s) they were gathered from.

We normalized the stimuli by peak amplitude using Audacity software (https://www.audacityteam.org). In one case (singing mouse) the stimuli were near the upper frequency threshold for sound detection in humans; to ensure our participants could hear them clearly, we pitch-shifted them down two octaves, also in Audacity.

All audio files can be accessed at http://github.com/themusiclab/animal-sounds.

### Experimental design

Participants listened to 16 pairs of stimuli (4 drawn at random with replacement from each of the four categories of frogs, birds, mammals, and insects), presented in random order. Because stimuli were selected at random with replacement, some participants received the same pair of stimuli more than once, providing an opportunity to measure intra-rater reliability.

On each trial, the first sound of a pair played after a half-second of silence. To maintain visual interest, as the sound played, an animal silhouette appeared on left side of the screen with a slight animated movement. The silhouette then disappeared, and after one second of silence, the second sound played, in conjunction with a second animal silhouette on the right side of the screen. The left stimulus always appeared first and the right stimulus always appeared second, but the order of presentation within pairs was randomized, such that the more-attractive sound preceded the less-attractive sound roughly as often as the reverse. Throughout, large text on the screen indicated the prompt “Which call do you **like** more?”. After one-tenth of a second of silence, both silhouettes reappeared and participants could respond. The means of response depended on device type: on desktop or laptop computers, the silhouettes were labelled with an “F” or “J” underneath (the left and right silhouette, respectively), and participants used their keyboard to enter their response; while on mobile devices (including tablets), participants tapped the left or right side of the screen. Participants on desktop or laptop computers could also enter responses by using a mouse click or touchscreen tap, but we encouraged them to use the keyboard to improve the validity of response time data.

Before and after the 16 trials described above, participants completed self-report questions on a variety of topics (see Demographic analyses, below).

Following the experiment, as an incentive for participation, we displayed a customized graphic indicating which of the four animal categories their responses agreed most with, along with descriptive information about the sounds of that animal category. This facilitated the sharing of results, spurring recruitment of more participants, a common strategy in gamified citizen science (*51*). Note that we did not indicate whether participants correctly identified the more-attractive sound during the experiment, precluding any learning effects during the experiment.

The experiment was implemented using the jsPsych ecosystem for behavioral experiments [https://jspsych.org; (*100*)] and hosted using the WorldWideLab backend (https://worldwidelab.org). Readers can try the experiment at https://themusiclab.org/quizzes/havoc. Code for the experiment is available at https://github.com/themusiclab/animal-sounds.

### Demographic analyses

We collected demographic data on participants to evaluate factors that could shape auditory preferences for non-human animal signals. Not all participants answered all demographic items; as such, we report the proportion of responses for each item as opposed to raw *N* s.

To identify “experts” in animal sounds, we asked “*Do you have a hobby or job where you identify animals by their sounds? For example birdwatching, herping, zookeeping, etc*.” Participants responding “*Yes*” were considered to be experts.

To identify “experts” in music, we asked “*Think of your skill at making music (using a musical instrument or singing). How would you rate your own skill?*”. Participants that answered “*I’m an expert*” or “*I have a lot of skill*” were considered to be experts, in contrast to those that answered “*I have no skill at all*”, “*I have some skill*”, or “*I’m a novice*” (i.e., this item was recoded as a binary category).

In addition to these data, which were used for planned analyses (see Main Text), we studied 10 additional variables in exploratory analyses of how experience and other variables impact participant responses.

These included:

- gender (male, female or other)
- age (in years) education (8 ordered responses from “*Some elementary/middle school*” to “*Completed graduate school*”)
- income (9 ordered responses from “*Under $10,000” to “Over $150,000*”), asked only of those reporting being located in the United States
- daily time spent making music (9 ordered responses from “*no time*” to “*> 4 hours*”)
- daily time spent listening to music (9 ordered responses from “*no time*” to “*> 4 hours*”)
- music lessons experience (binary variable)
- for participants who had completed music lessons, music skill in childhood, in relation to peers (5 ordered responses from “*They were a lot better than me*” to “*I was a lot better than them*”)
- music listening skill, with the text “*How good do you think your music listening skills are? (things like remembering melodies, hearing out of tune notes, or hearing a beat that is out of sync with the music)*” with responses on a sliding scale from “*I’m much worse than other people*” to “*I’m much better than other people*” with scale points unlabeled
- degree of music enjoyment, with the text “*In general, how much do you enjoy music?*” with a comparable sliding scale to (9), but anchors labeled “*I don’t enjoy music very much*” and “*I enjoy music very much*”

While many participants completed additional self-report items as part of their visit to https://themusiclab.org, we did not analyze them in this paper.

In a few cases, the variables required trimming or transformation. We detail these decisions below:

- Values for age were z-transformed.
- For the two sliding scales (music listening skill and degree of music enjoyment), we observed unusual response distributions with multiple peaks in responses, suggesting irregular response patterns. The default answer for both sliders was 500 (out of 1000); for both items, we omitted responses of exactly 500 (i.e., the participant did not move the slider: 587 out of 4165 participants and 229 out of 4164, respectively). We proceeded with two transformations:
  – For music listening skill we binarized the data using the midpoint of the scale, classifying only participants who moved the slider to the right as “skilled” relative to the other participants.
  – For degree of music enjoyment, we noted that about half the participants moved the slider all the way to the right anchor (1000), so rather than binarizing based on the midpoint of the scale, we classified those right-anchored participants as “enjoyers” relative to all other participants.

- While we used all data as described here in the statistical models (see below and in Main Text), to facilitate visual comparison of the effect sizes across each variable, in the figures, we binarized the numeric variables based on the mean value of the variable across participants.
- Finally, when including demographic covariates within our main model testing whether participants agreed with animals more than chance, we z-transformed the data.

We note that all trimming and transformation strategies for these exploratory analyses were determined without knowledge of the results.

All demographic variables were self-reported, thus will have some error based on biases in responses from participants. However, previous studies using self-disclosure of pitch perception or music expertise found strong correlations between the self-reported measures and performance on music perception tests (*101, 102*), suggesting that such items reliably track some aspects of cognition.

### Acoustic analyses

We measured 12 acoustic features using MATLAB. Three were calculated based on the waveform (dominant frequency, duration, and the amplitude modulation variance) and the remaining 9 were measured using `MIRtoolbox` (*103*), a package commonly used for music information retrieval: pitch (“best” prediction of the pitch), spectral centroid, brightness, roughness, Shannon entropy, attack slope, RMS energy (global energy), spectral skewness, and spectral spread. These metrics provide broad summaries of the sounds and do not describe acoustic patterning in detail.

Because the absolute level of each acoustic feature varies widely across species (e.g., pitch is far higher for *all* birds than for *all* frogs), we could not simply regress human preferences on acoustic characteristics. Instead, we first computed difference scores for each feature within each stimulus pair (i.e., subtracting the value for the less-attractive sound from the value for the more-attractive sound). These difference scores highlight what acoustic features are meaningful in distinguishing attractiveness across stimulus pairs, independently of the absolute level of specific acoustic features.

We studied the acoustic difference scores in relation to the estimated preferences of humans or animals. Specifically, we tested whether the more-attractive stimulus in each pair systematically varied as a function of acoustic features, relative to the less-attractive stimulus (e.g., for the feature of pitch, testing whether humans consistently selected the sound with lower pitch more often than expected by chance). Finally, we asked whether differences in acoustic features could predict whether humans chose the more-attractive stimulus (e.g., for the feature of pitch, testing whether larger differences in pitch between the two stimuli led to higher degrees of human-animal agreement).

In some cases, acoustic features could not be calculated (i.e., the function produced an NA value); we excluded those stimuli from analysis. This affected 5 stimulus pairs from the analysis of pitch (canary songs, likely due to the synthesized nature of the songs), 4 pairs from the analysis of spectral centroid (cricket and grasshopper songs, likely due to the broadband nature of their “clicks”), and 5 pairs from the analysis of attack slope (cricket and sparrow songs). Further, we assessed the distributions of difference scores for normality, and removed outliers: we omitted 3 pairs from the analysis of dominant frequency, 3 pairs from the analysis of spectral centroid, and 2 pairs from the analysis of roughness. Visual exclusion of outliers was done prior to conducting any statistical models and blind to the identity (e.g., species) of the outliers.

### Estimated strength of animal preference

For each stimulus pair, we used previous literature to generate an estimate of the strength of acoustic preference in the species that originally produced the sound. Given the heterogeneity of methods used in the studies from which we gathered the sounds, this required combining several approaches.

When studies had presented animals with binary choices between pairs of sounds, we computed the percent of animals choosing the preferred stimulus as the strength of preference (n.b., this measure is akin to the measure of human preference in our experiment, which was also a binary choice). When studies had used other behavioral measures, such as a count of the number of actions elicited by the sounds, we compared the magnitude of the behavioral result across stimulus types and calculated the percent of responses for the more-attractive sounds. When studies had used multiple behavioral responses with the same species and stimuli, we prioritized those that were most directly interpretable, e.g., those where animals’ movement toward or time spent near a speaker indicated their choice. We did so with the goal of enabling clear comparisons across studies and taxa.

For example, in a study of single phonotaxis tests in crickets, 78.3% of the crickets tested approached the preferred songs while only 19.1% of the crickets approached the unpreferred songs (*48*). Thus, out of all responses, 80% were for the preferred stimulus, indicating the strength of preference [78.3/(78.3 + 19.1) = 80.4]. In a more complex example, a study of swamp sparrow females found a mean of 6.9 copulation solicitation displays in response to more-attractive songs and only 3.9 displays in response to less-attractive songs (*39*). Thus, we estimated the strength of preference at 64% [6.9/(6.9 + 3.9) = 63.9].

While we typically used the exact values reported in published papers, when these values were not available, we used the PlotDigitizer tool (https://plotdigitizer.com) to estimate values visually from figures.

The strength of animal preference was variable (range: 55% – 93% preference), representing approximately 5:4 to 15:1 odds. We used this variation to test whether the strength of animal preference predicted the strength of human preference (see below). However, we also conducted analyses limited to relatively strong animal preferences. Specifically, we limited those analyses to stimuli with at least a 2:1 odds preference (≥ 66.67%; e.g., for every one individual that picked the less-preferred stimulus, two individuals picked the preferred stimulus) or at least a 3:1 odds preference (≥ 75%).

### Statistical approaches

To test whether participants selected the animal-preferred stimulus more than expected by chance, we used an intercept-only generalized linear mixed effects model (GLMM) with a binary response variable (1 = agreed with animals, i.e., chose the more-attractive sound, from the animal’s perspective; 0 = disagree) and a binomial link function, using `lme4` (*104*). We included Stimulus Pair ID, Species ID, and Participant ID as random effects, to license generalization across the sounds, species, and participants studied. These models all estimate the overall degree of agreement and test that estimate against a chance level of 0.5.

We used the same analytic approach for other key measures of interest, such as analyzing decision-making time, comparing the strength of preference between humans and animals, and assessing the influence of participant-level covariates on the main effects. Some data transformation was required for decision-making time analyses (i.e., square-root and z-transformed to improve model assumptions, limited to responses < 5 sec). To measure intra-rater reliability, we coded whether a participant selected the same (1) or different (0) sound across two repeated presentations, averaged these values for each stimulus pair, and tested this value against chance level of 50% with an intercept-only linear model (i.e., a one-sample *t*-test) across all stimuli.

For the acoustic feature analyses, we asked whether the value of any acoustic feature predicted whether humans or non-human animals preferred a stimulus in a pair, using mixed-effects models with an acoustic feature as the dependent variable, whether the stimulus was preferred (either by animals or by humans (greater than 50% human choice) in separate models) as an independent binary categorical factor, and Stimulus Pair ID and Species ID as random effects. We also asked whether the difference in a feature between the two stimuli could predict agreement between humans and animals, using linear mixed effects models with the size of the acoustic difference (computed by subtracting the value for the less-attractive sound from the value for the more-attractive sound) as the independent variable, average percent agreement (percent of human choices for the animal’s preferred stimulus) as the dependent variable, and Species ID as a random effect.

Full model specifications are available in the R Markdown manuscript available at https://github.com/themusiclab/animal-sounds. We used the car package for Type II Wald chi-square tests (*105*) for significance testing in the GLMMs and Type III analyses of variance with Satterthwaite’s method for the LMMs. When models failed to converge, we removed the random effect of species, and found nearly identical results. For any models with singular fits, the stimulus random intercept was dropped.

### Gender effects

While we limited our primary analyses to stimuli for which animals exhibited at least moderately strong preferences (2:1 odds, 66.67% preference), in preliminary analyses using the entire stimulus dataset, we found a significant effect of participant gender (male, female, or other) on their agreement with animals (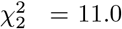, *p* < 0.01). Post-hoc contrasts indicated that male participants exhibited significantly greater agreement with other animals than female participants (*p* < 0.01), and no pairwise differences with participants with a gender of “other” (*p* > 0.61 for both). However, we note that the preferences of male and female participants were highly and significantly correlated (*r*^2^ = 0.7, *p* < 0.01), and as reported in the main text, the effect does not hold in our analysis limited to at least a 2:1 preference in animals. This indicates that the gender effect is primarily driven by responses to stimuli with the most ambiguity in animal preference (i.e., the stimuli where we have the least confidence in our estimation of the more-attractive stimulus).

Finally, we highlight that, in our primary analyses, all effects hold when limited to a single gender (significant difference from chance: male: *z* = 2.4, *p* = 0.02; female: *z* = 2.0, *p* = 0.04; other: *z* = 2.2, *p* = 0.03; correlation between animal preference and human preference strength: male: 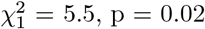, p = 0.02; female: 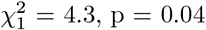, p = 0.04; other: 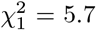, p = 0.02). As such, we suggest caution in interpreting gender effects and encourage future research on this subject.

## Supplementary tables and figures

**Table S1.**
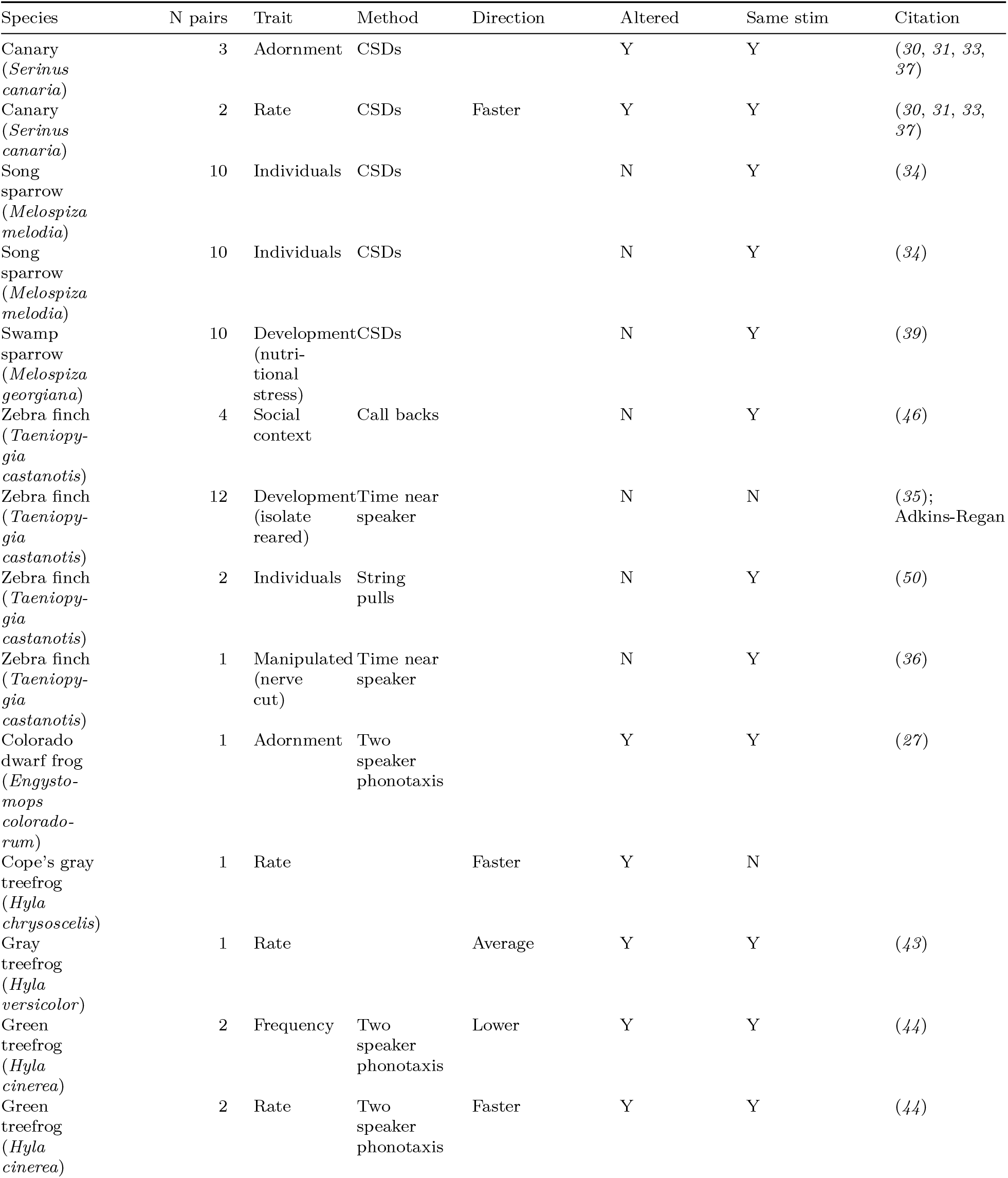
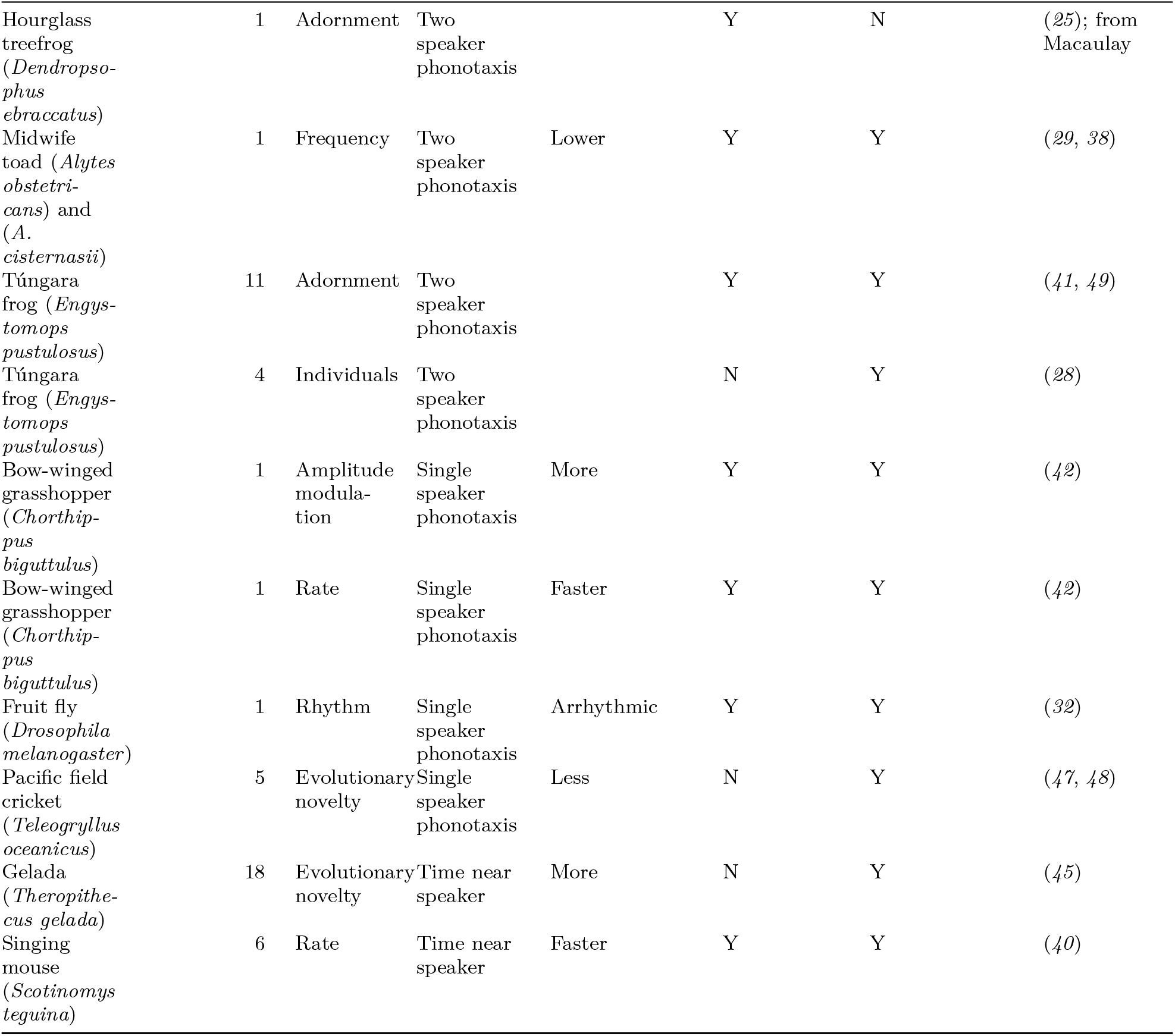
Description of stimuli. Note. “N” refers to the number of stimulus pairs; “Altered” refers to whether the sounds were experimentally manipulated; “Same stim” refers to whether or not we were able to obtain and use the exact stimuli tested on the animals themselves, or used other sources; “CSDs” refers to copulation solicitation displays.

**Table S2.** Acoustic results. Note. “DF” refers to degrees of freedom. A “Diff” comparison asks whether the difference in a feature between the stimuli in the pair predicted agreement between humans and non-human animals; “Animals” and “Humans” comparisons ask whether that group demonstrates a directional preference for the feature. “AM var” refers to amplitude modulation variance and “RMS energy” refers to the root-mean-square energy (overall measure of energy in the sound).

| feature | comparison | DFnum | DFden | F | p | direction |
| --- | --- | --- | --- | --- | --- | --- |
| Dominant frequency | Diff | 1 | 61.1 | 0.1 | 0.75 |  |
| Dominant frequency | Animals | 1 | 66.0 | 3.5 | 0.07 |  |
| Dominant frequency | Humans | 1 | 66.0 | 0.2 | 0.65 |  |
| Duration | Diff | 1 | 64.5 | 0.4 | 0.54 |  |
| Duration | Animals | 1 | 69.0 | 5.4 | **0.02** | Prefer longer |
| Duration | Humans | 1 | 69.0 | 0.8 | 0.37 |  |
| Pitch | Diff | 1 | 59.3 | 12.2 | **< 0.01** |  |
| Pitch | Animals | 1 | 64.0 | 0.1 | 0.78 |  |
| Pitch | Humans | 1 | 64.0 | 11.4 | **< 0.01** | Prefer lower |
| AM var | Diff | 1 | 66.1 | 0.4 | 0.54 |  |
| AM var | Animals | 1 | 127.8 | 0.2 | 0.66 |  |
| AM var | Humans | 1 | 127.8 | 0.1 | 0.71 |  |
| Spectral centroid | Diff | 1 | 60.9 | 2.8 | 0.10 |  |
| Spectral centroid | Animals | 1 | 62.0 | 6.1 | **0.02** | Prefer higher |
| Spectral centroid | Humans | 1 | 62.0 | 2.6 | 0.11 |  |
| Skew | Diff | 1 | 44.7 | 1.4 | 0.24 |  |
| Skew | Animals | 1 | 128.0 | 1.9 | 0.17 |  |
| Skew | Humans | 1 | 128.0 | 1.3 | 0.26 |  |
| Spread | Diff | 1 | 29.2 | 1.2 | 0.29 |  |
| Spread | Animals | 1 | 69.0 | 2.7 | 0.10 |  |
| Spread | Humans | 1 | 69.0 | 3.9 | 0.05 |  |
| Shannon entropy | Diff | 1 | 31.3 | 1.3 | 0.26 |  |
| Shannon entropy | Animals | 1 | 127.8 | 0.2 | 0.67 |  |
| Shannon entropy | Humans | 1 | 127.8 | 1.7 | 0.19 |  |
| Roughness | Diff | 1 | 65.8 | 0.6 | 0.46 |  |
| Roughness | Animals | 1 | 125.0 | 0.1 | 0.72 |  |
| Roughness | Humans | 1 | 125.0 | 0.9 | 0.35 |  |
| Attack slope | Diff | 1 | 63.0 | 1.2 | 0.28 |  |
| Attack slope | Animals | 1 | 64.0 | 0.5 | 0.48 |  |
| Attack slope | Humans | 1 | 64.0 | 0.2 | 0.63 |  |
| RMS energy | Diff | 1 | 65.3 | 0.2 | 0.67 |  |
| RMS energy | Animals | 1 | 128.1 | 0.1 | 0.73 |  |
| RMS energy | Humans | 1 | 128.1 | 1.1 | 0.30 |  |
| Brightness | Diff | 1 | 54.8 | 0.9 | 0.34 |  |
| Brightness | Animals | 1 | 128.0 | 8.3 | **< 0.01** | Prefer brighter |
| Brightness | Humans | 1 | 128.0 | 1.6 | 0.21 |  |

**Table S3.** Statistical results for demographics. Note: The chi-squared tests are omnibus tests from the GLMMs described under “Demographic analyses” in the Methods.

| trait | Chisq | DF | p |
| --- | --- | --- | --- |
| animal sounds expert | 1.9 | 1 | 0.17 |
| music expert | 1.3 | 1 | 0.25 |
| music lessons | 0.4 | 1 | 0.51 |
| ability wrt peers | 3.1 | 1 | 0.08 |
| time making music | 0.7 | 1 | 0.41 |
| time listening to music | 8.5 | 1 | **< 0.01** |
| listening skill | 1.5 | 1 | 0.22 |
| music enjoyment | 0.3 | 1 | 0.56 |
| education | 1.3 | 1 | 0.26 |
| age | 0.7 | 1 | 0.42 |
| income | 1.8 | 1 | 0.18 |
| gender | 3.1 | 2 | 0.21 |

**Fig. S1.**
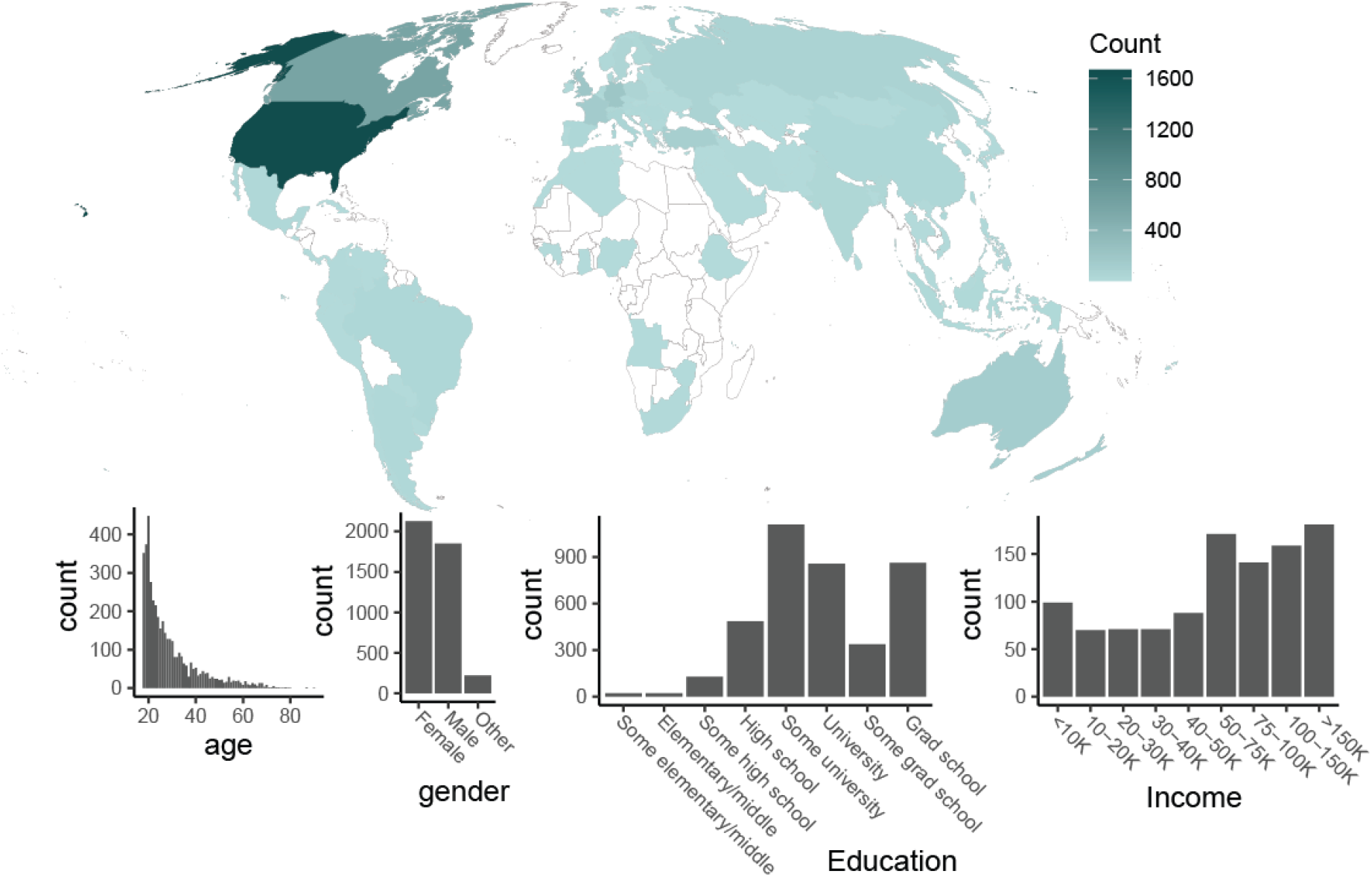
Participant demographics. The map depicts the distribution of participants, with darker-shades indicating a larger number of participants who were resident in each country. The bar plots show the distributions of participants’ age (in years), gender, education, and income levels of participants.

**Fig. S2.**
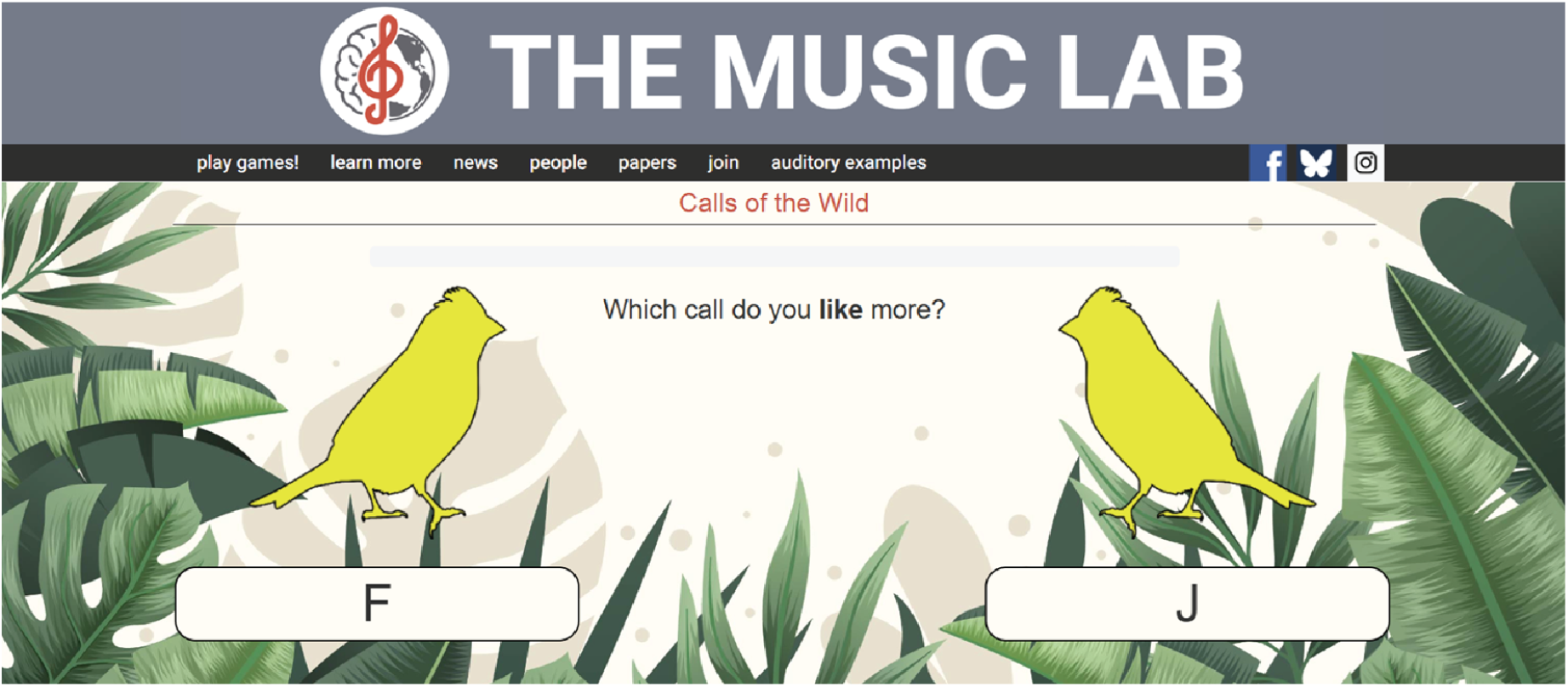
User interface. The screenshot shows the experiment’s user interface, as appearing for participants who used a desktop or laptop computer. This screen appears after both sounds on a given trial have played and the participant responds by pressing the F or J key on their keyboard (or clicking or tapping one of the two bird silhouettes).

**Fig. S3.**
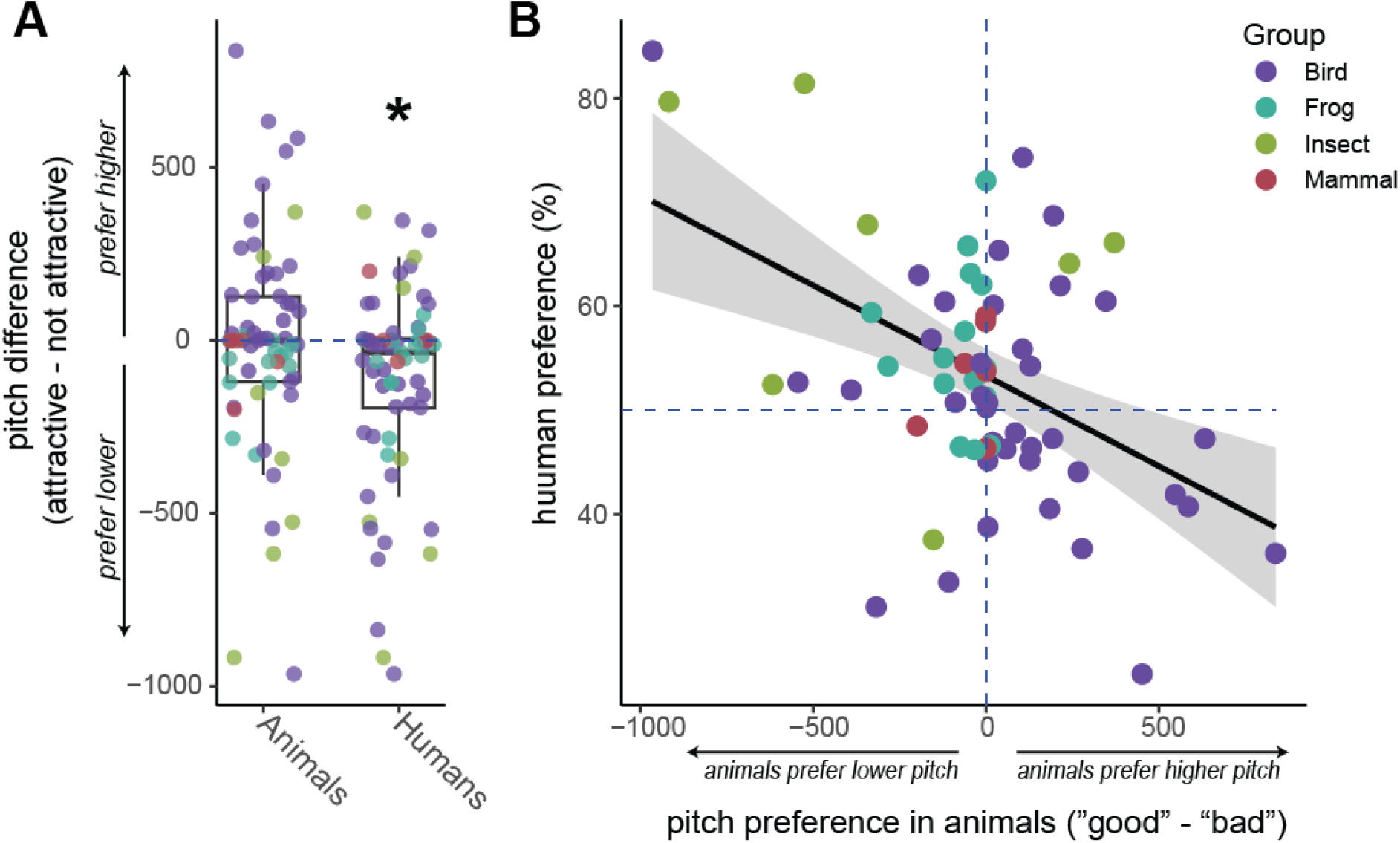
Influence of pitch on animal and human preferences. (**A**) Across all stimulus pairs, humans tended to choose the stimulus with the lower pitch, while there was no consistent preference across the other animals. (**B**) The difference in pitch within a pair of stimuli was predictive of the degree of human agreement with animals (“human preference”; i.e., humans were more likely to agree with animals when the animal-preferred stimulus was lower in pitch).

